# The contraction-expansion behaviour in the demosponge *Tethya wilhelma* is diurnal and light-controlled

**DOI:** 10.1101/2022.07.06.499042

**Authors:** Sarah B. Flensburg, Anders Garm, Peter Funch

## Abstract

Sponges (phylum Porifera) are metazoans without muscles and nervous system. Still, they perform coordinated behaviours, such as whole body contrations. Previous studies have indicated diurnal variability in number of contractions, and in expression of circadian clock genes. Here we show that diurnal patterns are present in the contraction-expansion behaviour of the demosponge *Tethya wilhelma* using infrared videography and a simulated night/day-cycle including sunset and sunrise mimic. In addition, we show that this behaviour is at least strongly influenced by the ambient light intensity and therefore implicates light-sensing capabilities in this sponge species. This is backed by our finding that *T. wilhelma* consistently contracts at sunrise, and that this pattern disappears both when the sponge is kept in constant darkness and when in constant light.

## Introduction

Sponges (phylum Porifera) are aquatic animals with a world-wide distribution found at all water depths. The group comprises over 20,000 described species across four extant classes of which the class Demospongiae contains most of the species and is the most abundant (De Voogd et al., 2021; Van Soest et al., 2012). The sponge body plan is fundamentally organized around cavities and canals where water flows through the sponge (Asadzadeh et al., 2020). Despite their relatively simple body organization, sponges possess a number of different cell types but lack muscle and nerve cells (Lavrov and Kosevich, 2014; Musser et al., 2021). Still, they are able to perform coordinated behaviours such as whole-body contractions, and some sponges even have the ability to move (Bond, 2013; Nickel, 2006). Contractions can happen spontaneously. They can also be initiated by a range of stimuli including different chemicals (Elliott and Leys, 2010; Ellwanger et al., 2007; Ellwanger and Nickel, 2006; Goldstein et al., 2020), or by a mechanical stimulus locally on the exopinacoderm and spread as a contraction wave across the surface of the sponge (Elliott and Leys, 2007; Nickel et al., 2011). Also, shaking of the freshwater sponge *Ephydatia muelleri* causes a contraction, and exposure to high loads of ink particles causes it to contract and expel waste particles from its water canal system (Elliott and Leys, 2007). The mechanism initiating this behaviour has not been identified but primary cilia located in the oscular area have been suggested to function as mechanoreceptors involved in water flow regulation and sponge body contractions (Ludeman et al., 2014). Despite the lack of neurons, stimulation of one part of the sponge often result in changes in behaviour throughout the entire sponge, suggesting coordination, possibly facilitated by chemical messengers (Kealy et al., 2019). Light also influences sponge behaviours, including the release of larvae, which in some species is correlated with the lunar cycle (Amano, 1986; Amano, 1988; Hoppe and Reichert, 1987; Neely and Butler, 2020). Some sponge larvae likewise respond to changes in light conditions, suggesting that they possess a light-sensory system (Leys et al., 2002). However, genetic data suggest that sponges lack the photopigment opsin, most often involved in animal vision, and evidence from the demosponge *Amphimedon queenslandica* indicates that the larvae here use cryptochromes instead (Mah and Leys, 2017; Rivera et al., 2012). In addition to performing behaviours normally associated with a nervous system, sponges also possess many of the genes associated with the nervous system in other animals. This has lead to the hypothesis that sponges, while not having a nervous system *per se*, might have a kind of pre-nervous system able to perform some of the functions of a fully developed nervous system (Leys, 2015).

Diurnal activity patterns are commonly observed in many different types of organisms. In animals, a typical pattern consists of alternating periods of inactivity, sleep, which can be observed across a wide range of species (Siegel, 2008), including animals without a conventional brain such as the cnidarian polyp *Hydra vulgaris* (Kanaya et al., 2020) and the jellyfish *Cassiopea* sp. (Nath et al., 2017). Studies of diurnal rythms in animals without a nervous system are very limited. In a long term *in situ* study Reiswig (1971) showed a synchronized diurnal cycle of body contractions and expansions of the demosponge *Tectitethya crypta*. Another demosponge *Tethya wilhelma* also seems to exhibit diurnal variations in body contractions, as indicated by the results from two specimens tested under laboratory conditions with an artifical 12h:12h light:dark cycle (Nickel, 2004). The control mechanisms behind the diurnal rhythm were not tested, but molecular work from the demosponge *Amphimedon queenslandica* revealed a diurnal expression of two metazoan circadian clock genes suggesting that it could be at least partly intrinsically regulated (Jindrich et al., 2017). Not surprisingly there seems to be a lot of variation between species and an *in situ* study on the demosponge *Aplysina aerophoba* did not find any diurnal rhythms in pumping activity (Pfannkuchen et al., 2009).

In this study we have assessed the indicated diurnal rhythm in the body contraction-expansions of the sponge *T. wilhelma*, and tested the potential influence of the ambient light intensity. We confirmed the diurnal rhythms in the contractions and through manipulations of the light regime we also found evidence for the rhythm to at least largely be controlled extrinsically by the light level.

## Materials and methods

### Culturing of sponges

Specimens of *Tethya wilhelma* Sarà et al., 2001 were collected at the National Aquarium of Denmark, the Blue Planet, (Copenhagen, Denmark) in a coral reef display tank with sea water of 24°C and 33 psu and transported to Aarhus University. The specimens had a size of 0.5-1.0 cm in diameter. After arrival, they were transferred to individual wells of a six well culture plate with a 2.5 cm diameter black plastic disk placed at the bottom of each well for sponge attachment. The well plate was then placed in a 25-litre glass tank with seawater collected at Lemvig, Denmark, adjusted to 32-34 psu using artificial sea salt (Instant Ocean, Aquarium Systems, France). The water temperature was kept at 22°C controlled by the room temperature. The water was filtered and circulated using an EHEIM eXperience 150 external filter pump incl. a biological filter (EHEIM GmbH & Co. KG, Germany). Light was supplied with a Fluval Plant Nano Bluetooth LED light (Rolf C. Hagen Inc., Canada), which was programmed to simulate sunrise from 6:00 AM to 7:00 AM, daylight from 7:00 AM to 5:00 PM, sunset from 5:00 PM to 6:00 PM, and dark from 6:00 PM to 6:00 AM. This corresponds to a light regime of 12 hours of complete darkness, and 12 hours of varying light. The light pre-set “Tropical River” was used at 1/10 of the original light intensity to prevent algal growth. Approximately one third of the water in the tank was exchanged every three weeks throughout the experimental period.

### Experimental setup

The sponges were transferred to a submerged clear acrylic box (12/17/4.8 cm, H/W/D) inside the holding tank. Light could enter directly from above and from the sides but the back was covered with black fabric to provide a uniform dark background for video recordings. A weight was placed on top of the box to make sure it wasn’t moved by the water currents in the tank during the video recordings. The sponges were video recorded from the outside of the tank to document changes in the body sponge volume measured as changes in the projected area. Recordings were made with a Panasonic HC-VX980 digital camcorder (Panasonic Corp., Japan) using the night mode setting. Two infrared LED light sources (peak emission at 940 nm) were used for illumination during the night (Fig. S1).

### Test of diurnal rhythm

To characterize the contraction pattern of *Tethya wilhelma* under simulated natural dark/light conditions, six specimens were recorded for 72 hours each. One at a time, the sponges were transferred to the recording chamber at least three hours before the start of each recording session to mimimize the influence of the handling. All recordings were started at 6:00 PM on the first day and after 72 hours of recording, the sponge was transferred from the recording chamber and back into the well plate. The influence of the light regime on contraction cycles in the sponge *T. wilhelma* was investigated by subjecting the six specimens to 72 hours of constant light and 72 hours of constant dark. These two treatments were separated by a recovery period with a 12:12 hour dark:light cycle for seven days. During the recovery period, the sponges were placed in another tank. Three sponges were tested at the same time and they were again transferred to the recording chamber at least three hours before the start of the experiment. Because the experimental period started at 6:00 PM, a simulated sunset would be included at the onset of the constant darkness experiments but not the constant light experiment. To ensure complete control of the light environment in the experimental tank it was covered by a light proof box also including the camera and light sources.

### Processing of video material

Single frames were grabbed from the videos with a time resolution of 100 s and transferred to the software program ImageJ (Rasband, 2020) as an image stack, and a scale bar was added using the plate the sponges were attached to as a reference. The images were converted into 8-bit greyscale, cropped, and had the brightness and contrast adjusted to optimize the contrast between sponge and background. Based on the difference in contrast between the bright sponge and the dark background, the part of the image containing the sponge could be selected by converting the image to a binary image by setting an appropriate threshold and applying it to the image stack. The projected area of the sponge was then measured using a “Measure Stack” macro in ImageJ.

### Analysis of contraction patterns

The area measuremensts of each sponge were normalized by the maximum area during the 72-hour period to obtain a relative projected area (RPA) (see Goldstein et al. (2020)). The mean RPA for each 12-hour period (6:00 AM to 6:00 PM and 6:00 PM to 6:00 AM) was calculated and the number of fully contracted states, indentified as RPA minima, were counted for each 12-hour time period as well. The mean contraction amplitude was calculated for each 12-hour time period as the mean change in RPA between the initiation of the contractions and the fully contracted states during that time period. The mean contraction area was calculated as the mean RPA of the fully contracted states during each 12-hour period. Lastly, the minimum and maximum RPA during each 12-hour time period was identified. If a contraction spanned both dark and light period then it was assigned to the period where the contraction was initiated. The sponges are constantly making small changes of their volume and accordingly a threshold of minimum 7% reduction in RPA was set before it was considered a contraction. This threshold was choosen, as contractions with a greater than 7% change in RPA displayed the typical shape of a contraction, with a rapid contraction, followed by a slower expansion phase. A combined graph was produced for each of the three experiments, combing the individal specimens (N=5/6) and the individual days (N=3) into one graph to reveal possible consistent patterns in the timing of the individual contractions. Here 95% confidence intervals were reported.

### Statistical analyses

The statistical analyses were performed in RStudio version 2022.02.1 (RStudio Team, 2022), using the R packages “lme4” (Bates et al., 2015) and “lmerTest” (Kuznetsova et al., 2017). Plots were made using the package “ggplot2” (Wickham, 2016) and Microsoft Excel. Linear mixed-effect models (LMEM’s) were fitted, that met the assumptions of normal distribution, homoscedasticity, and independence of residuals. As response variables, mean RPA, mean contraction amplitude, mean RPA of contractions, minimum RPA, and maximum RPA were used. As explanatory variable, day period (either day or night) was added as a fixed factor, and sponge ID and day number were added as random effects. For the analysis of the number of fully contracted states, a generalized linear mixed model (GLMM) with poisson distribution was fitted. This model met the same assumptions and contained the same explanatory variables and random effects as the LMEMs. In the comparisons between experiments (dark/light, constant dark, and constant light), experiment was added as a fixed factor instead of period, and *post-hoc* tests were performed by multiple comparisons of means using Tukey contrasts and single-step method for adjusting *p-*values using the ‘glht’-function from the R package “multcomp” (Hothorn et al., 2008). The critical p-value was set to 0.05 for all tests. The standard deviations (± s.d.) are calculated based on number of specimens with the experimental days pooled together.

## Results

### Contraction behaviour under normal conditions and constant dark/light

One sample had to be excluded from the analysis in the constant dark experiment because the sponge moved out of the focus plane of the scale bar. The sample sizes were therefore as follows; dark/light experiment (N=6), constant dark experiment (N=5), and constant light experiment (N=6).

The summarized contractions-expansion diagrams showed a pattern of one contraction happening consistently shortly after the time of sunrise in the simulated dark/light-experiment (Fig. 1A, Fig. S2). This pattern was not observed in neither the constant dark, nor the constant light experiment (Fig. 1B, C, Fig. S3, S4).

**Figure 1.**
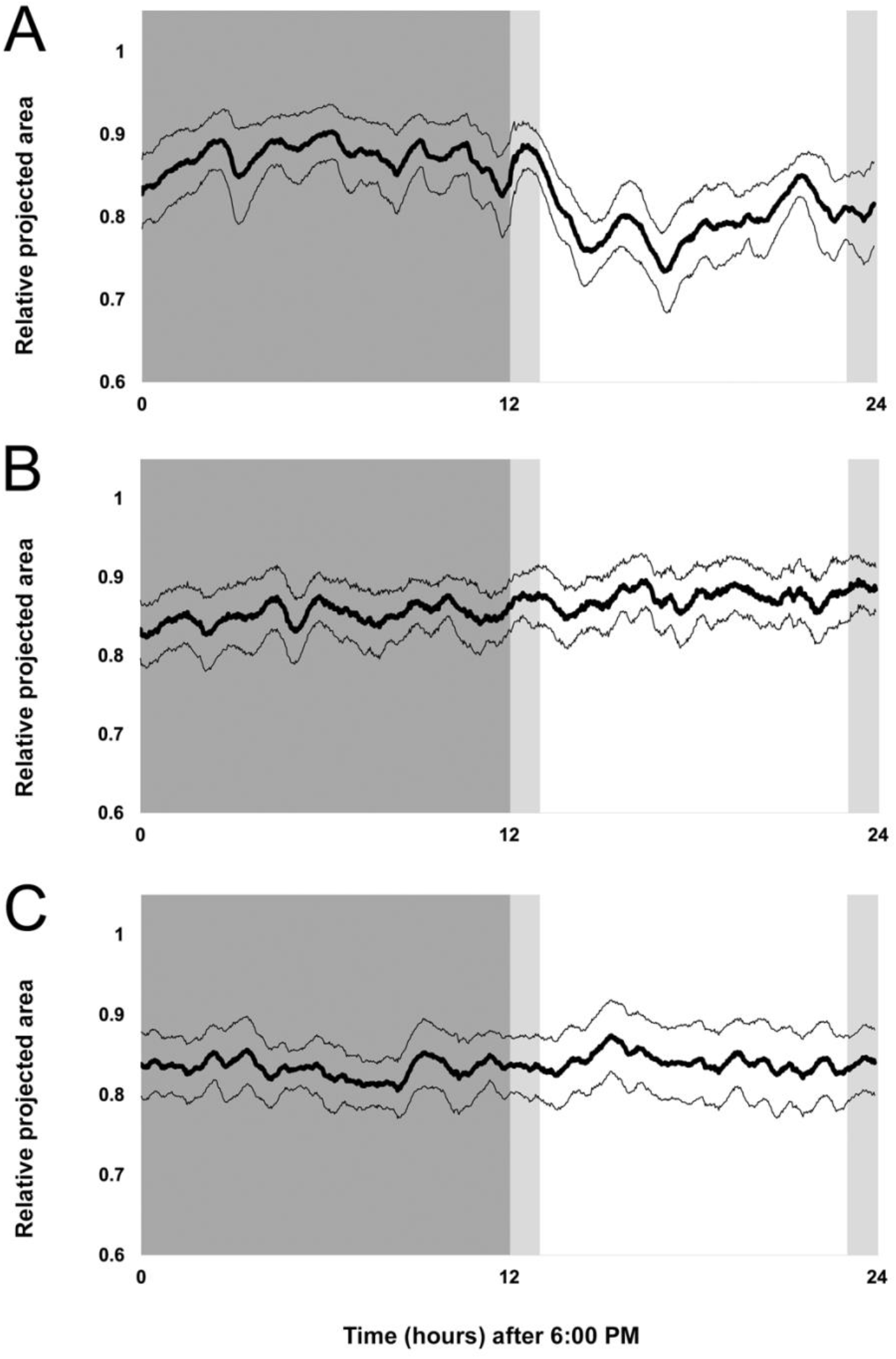
Summarized contraction-expansion graphs of specimens of *T. wilhelma*. Mean RPA over three days for all specimens are shown with thick lines and 95% confidence interval with thin lines. A) Simulated night/day-cycle (N=6), B) constant dark (N=5) and C) constant light (N=6). Dark grey indicates night period, light grey indicates periods with sunrise/sunset.

When sponges were kept under normal light conditions, the mean RPA was 0.873 (±0.026) for the night periods and 0.803 (±0.050) for the day periods which is a significant difference (LMEM, *t*=3.6, *p*=0.0011) (Fig. 2A). The mean number of fully contracted states also differed significantly between night and day periods (LMEM, *z*=−2.4, *p*=0.016) and the mean was 2.22 (±0.78) during night periods and 3.61 (±0.77) during day periods (Fig. 2B). The relative contraction aplitude was 0.214 (±0.076) during the night and 0.190 (±0.058) during the day (Fig. 2C). This difference was not significant (LMEM, *t*=1.2, *p*=0.239). The mean contraction RPA was also not significantly different at night compared to daytime, 0.705 (±0.073) and 0.676 (±0.083) respectively (LMEM, *t*=1.6, *p*=0.135) (Fig. 2D). The mean minimum RPA was 0.702 (±0.062) during the night period and 0.645 (±0.087) during the day period respectively, which is a significant difference (LMEM, *t*=2.3, *p*=0.028) (Fig. 2E). Finally, the maximum RPA did not differ between night period, 0.941 (±0.032), and day period, 0.933 (±0.020) (LMEM, *t*=0.4, *p*=0.684) (Fig. 2F). Under constant darkness and constant light conditions, no significant differences were found between night period and day period in any of the examined parameters (Table 1, Fig. 2A-F).

**Table 1.** Results from constant dark and constant light experiment in *T. wilhelma*. Mean (± s.d.) values for each of the six examined parameters during the night periods and the day periods. For the comparisons between periods, *t*- and *p*-values are presented, exept for no. fully contracted states, where *z*- and *p*-values are presented.

| **Constant darkness experiment** |  |  |  |  |
| --- | --- | --- | --- | --- |
|  | **Night period** | **Day period** | ***t* (*z*)** | ***p*** |
| Mean RPA | 0.852 ($\pm 0.028$) | 0.875 ($\pm 0.028$) | -1.5 | 0.155 |
| No. Fully contracted states | 3.8 ($\pm 0.61$) | 3.93 ($\pm 0.86$) | -0.2 | 0.853 |
| Rel. contraction amplitude | 0.135 ($\pm 0.010$) | 0.150 ($\pm 0.020$) | -1.0 | 0.314 |
| Mean contraction RPA | 0.755 ($\pm 0.032$) | 0.767 ($\pm 0.042$) | -1.0 | 0.350 |
| Min. RPA | 0.723 ($\pm 0.039$) | 0.748 ($\pm 0.035$) | -1.6 | 0.131 |
| Max. RPA | 0.926 ($\pm 0.024$) | 0.946 ($\pm 0.024$) | -1.1 | 0.280 |
| **Constant light experiment** |  |  |  |  |
|  | **Night period** | **Day period** | ***t* (*z*)** | ***p*** |
| Mean RPA | 0.833 ($\pm 0.041$) | 0.841 ($\pm 0.055$) | -0.5 | 0.652 |
| No. Fully contracted states | 3.39 ($\pm 0.49$) | 3.5 ($\pm 0.78$) | -0.2 | 0.857 |
| Rel. contraction amplitude | 0.169 ($\pm 0.017$) | 0.168 ($\pm 0.013$) | 0.03 | 0.977 |
| Mean contraction RPA | 0.725 ($\pm 0.043$) | 0.734 ($\pm 0.046$) | -0.7 | 0.485 |
| Min. RPA | 0.708 ($\pm 0.037$) | 0.705 ($\pm 0.040$) | 0.3 | 0.746 |
| Max. RPA | 0.931 ($\pm 0.037$) | 0.939 ($\pm 0.028$) | -0.4 | 0.668 |

**Figure 2.**
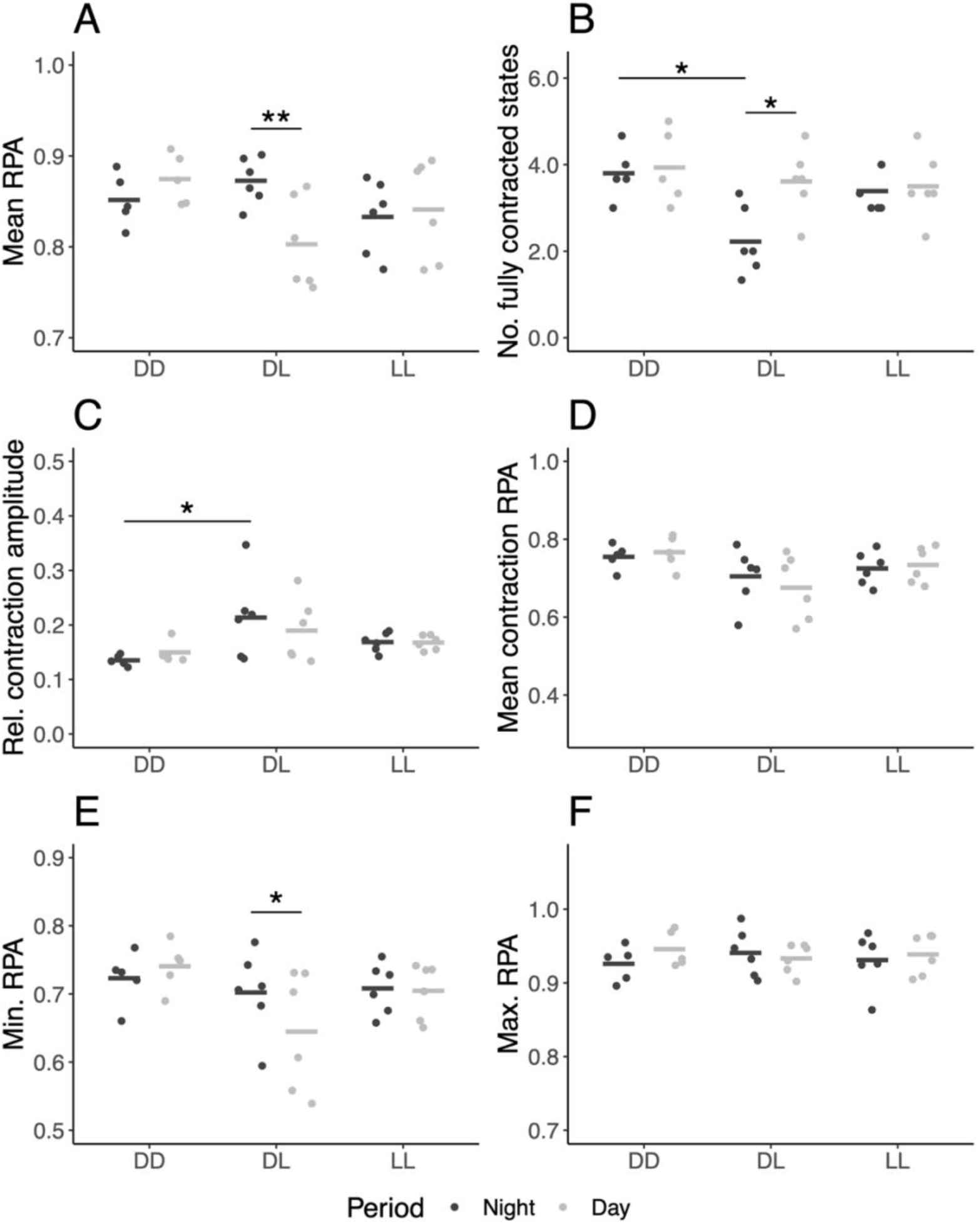
Influence of light regime on contraction behaviour of *T. wilhelma*. Differences between experiments, and between night period (dark grey) and light period (light grey) in mean RPA (A), number of fully contracted states (B), relative contraction amplitude (C), mean contraction RPA (D), minimum RPA (E), and maximum RPA (F). Asterisks indicate a significant difference; *=p<0.05, **=p<0.01. Points are individual specimens (N=6 for DL and DD, N=5 for LL), bars are means. DD = constant dark, DL = dark/light, and LL = constant light.

### Normal light regime vs constant dark/light

When comparing the timewise day periods between the experiments, the mean RPA was significantly higher in the constant dark experiment compared to the dark/light according to the linear mixed-effect model (LMEM, *t*=−2.3, *p*=0.040), but in the multiple comparisons test it is only a trend (Tukey comparisons *post hoc*: *p*=0.054) (Fig. 2A). Also, minimum RPA was significantly higher in the constant dark experiment compared to the dark/light experiment according to the linear mixed-effect model (LMEM, *t*=−2.3, *p*=0.047), but again only a trend according to the multiple comparisons test (Tukey comparisons *post hoc*: *p*=0.055) (Fig. 2E). When comparing the timewise night periods between the three experiments, the number of fully contracted states was significantly higher in constant dark experiment compared to the dark/light experiment (Tukey comparisons *post hoc*: *p*<0.025) (Fig. 2B), and the mean relative contraction amplitude was significantly lower in the constant dark experiment compared to the dark/light experiment (Tukey comparisons *post hoc*: *p*=0.028) (Fig. 2C). The constant light experiment did not differ significantly from the light/dark experiment for any of the parameters (Fig. 2A-F).

## Discussion

The present study shows a diurnal rhythm in the contraction patterns of *Tethya wilhelma*. Specifically when comparing night and day periods, the mean RPA was significantly higher during the night (Fig. 2A), the number of fully contracted states was significantly higher during the day (Fig. 2B), and finally, the minimum RPA measurements during each 12-hour time period was significantly higher during the night (Fig. 2E). These differences suggest that *T. wilhelma* is more expanded during the night, and is more actively contracting during the day. The constant light and constant dark experiments on *T. wilhelma* showed, that any significant differences found in the dark/light experiment disappeared under these constant light regimes. In complete darkness the number of fully contracted states is similar to daytime in the dark/light experiment and significantly higher than at nighttime indicating that prolonged darkness increases the frequency of contractions. In addition, the amplitude of the contractions during the night period in the constant darkness experiment is significantly smaller than the amplitude of the contractions during nighttime under normal light conditions, showing that in addition to the contractions happening more frequently, they are also smaller (Fig. 2C). The multiple comparisons *post-hoc*-tests failed to show differences among experiments in mean RPA and minimum RPA during the timewise dayperiods. This is likely due to low statistical power caused by unequal sample sizes between experiments, and as the differences in means show, effects sizes are clearly large and confirm that the sponges are more expanded during darkness, than during light (Fig. 2A, E).

The finding from the present study, that *T. wilhelma* has a diurnal variation in contraction patterns is in line with another study on this species by Nickel (2004), where projected area measurements were also used to count and compare the number of contractions during day and night. However, the study by Nickel (2004) was limited to two specimens, so the present study is the first to statistically prove this diurnal rhythm. In addition, our results show that this diurnal rhythm is not restricted to the number of contractions alone but is also present in two additional parameters; minimum and mean relative projected area. It should be noted, though, that these three parameters are not completely independent, since more contractions equal more minima, and more time spent in a contracted state leads to lower mean RPA. For functional considerations of the contractions in sponges it is important that although contractions result in changes of the overall volume of the sponge, these changes mainly concern the volume of the water canal system (Nickel et al., 2011). Sponge contractions have been suggested to serve different functions, including servering as a cleaning mechanism to expel unwanted particles by flusing the water canal system (Elliott and Leys, 2007; Nickel, 2004). In this case diurnal periodicity in contraction-expansion behaviour could be related to maintenance of the water canal system. Further studies are needed to reveal the relationship between diurnal activity patterns and individual elements of the water canal system and to test the hypothesis that *T. wilhelma* is more actively filtering during daytime and thus needs more flushing in this period.

### Control of the diurnal rhythm

Since our study shows the absence of diurnal rhythms in *T. wilhelma* under constant dark and constant light conditions, it implicates that contraction behaviour is, at least in part, regulated exogenously by the ambient light intensity, and as a consequence, that this sponge has light sensing capabilities. Nickel (2004) used an external flash to capture images of the contraction states of *T. wilhelma*. Futher studies are needed to determine a possible behavioural impact of exposure to short flashes of light during the night, and clarify if this could influence the natural contraction patterns by, for instance, increasing the frequency of contractions. In our study we used an infrared lightsource to avoid this possible impact. Also, our use of a light capable of simulating sunrise/sunset results in more natural changes of light levels. Indeed, we found a consistent pattern of a contraction happening shortly after simulated sunrise. This behaviour was not reported by Nickel (2004) who used an artificial 12:12 light:dark cycle. This further strengthens our hypothesis, that *T. wilhelma* has light sensing capabilities, and that sunrise could serve as a ‘guide’ for when to increase activity i.e. contractions, potentially flushing the canal system to be ready for increased daytime filtration.

The diurnal rhythm in *T. wilhelma* is in line with genomic studies on another demosponge *Amphimedon queenslandica*. Here, diurnal variations were found in expression of two circadian clock genes, but the variations became less pronounced when the sponge was subjected to constant darkness (Jindrich et al., 2017). This suggest at least some endogenous control of diurnal rhythms alongside the light control shown by our results. Because many sponge species have a diverse microbiome including photosynthesizing symbionts (Bickmeyer et al., 2019; Thomas et al., 2016), the diurnal rhythm of the sponge could be influenced by this microbiome, and decreasing levels of oxygen inside the sponge at night could account for the diurnal rhythm in behaviour. This however is contradicted by the very similar patterns observed under constant darkness and constant light. Light seems to play a similar role in the cnidarians *Hydra vulgaris, Tripedalia cystophora, and Copula sivickisi* that all show diurnal rhythms in behaviour. Also, the diurnal activity patterns disappeared under either constant light or constant dark conditions in *H. vulgaris* (Garm et al., 2012; Kanaya et al., 2020). Changes in behaviour in response to light is also seen among the sistergroup to animals, the choanoflagellates. Brunet et al. (2019) revealed that colonies of the choanoflagellate *Choanoeca flexa* inverted when subjected to changing light levels, showing that even simple holozoans are capable of modulating their behaviour in response to light, demonstrating that such behaviour in sponges is not at all unexpected.

## Acknowledgements

We are grateful to the National Aquarium of Denmark, the Blue Planet, for providing animals for our study, and to Dorthe Due Theilade for transport to our laboratory. In addition, we are grateful to Alexandra B. Flensburg for help with designing the experimental setup, and to Josephine Goldstein and Clémence Rose for helping with the statistical analyses.

## Author contributions

Conceptualization: S.B.F, A.G., P.F.; Methodology: S.B.F, A.G., P.F.; Formal analysis: S.B.F.; Investigation: S.B.F.; Resources: A.G., P.F.; Writing - original draft: S.B.F.; Writing - review and editing: S.B.F, A.G., P.F.; Visualization: S.B.F.; Supervision: A.G, P.F.; Funding acquisition: P.F.

