## Supplementary material for "The contraction-expansion behaviour in the demosponge *Tethya wilhelma* is diurnal and light-controlled"

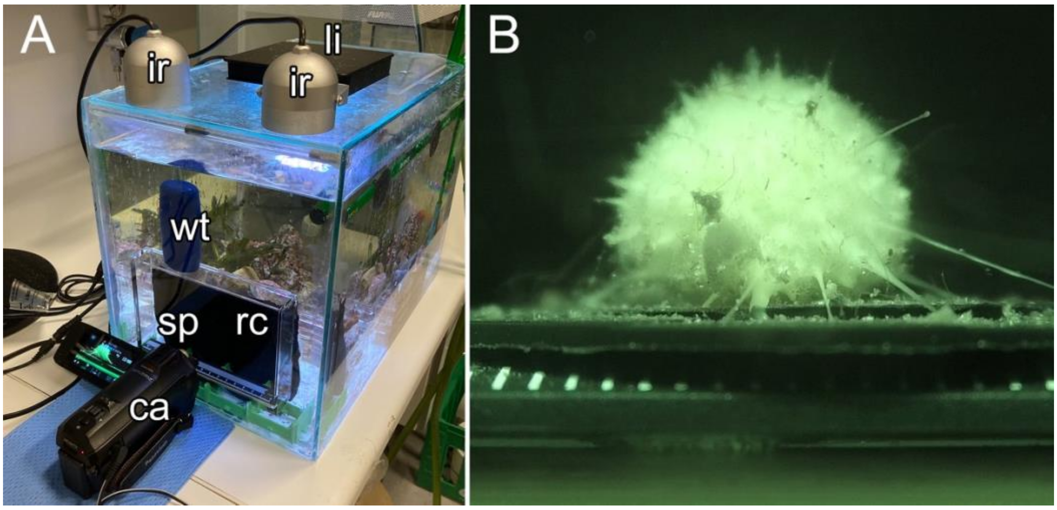


**Figure S1. Experimental setup.** Tank with recording equipment (A). Camera view of sponge during recording (B). ca = camera, sp = sponge, rc = recording chamber, wt = weight, ir = infrared light sources, li = aquarium light.

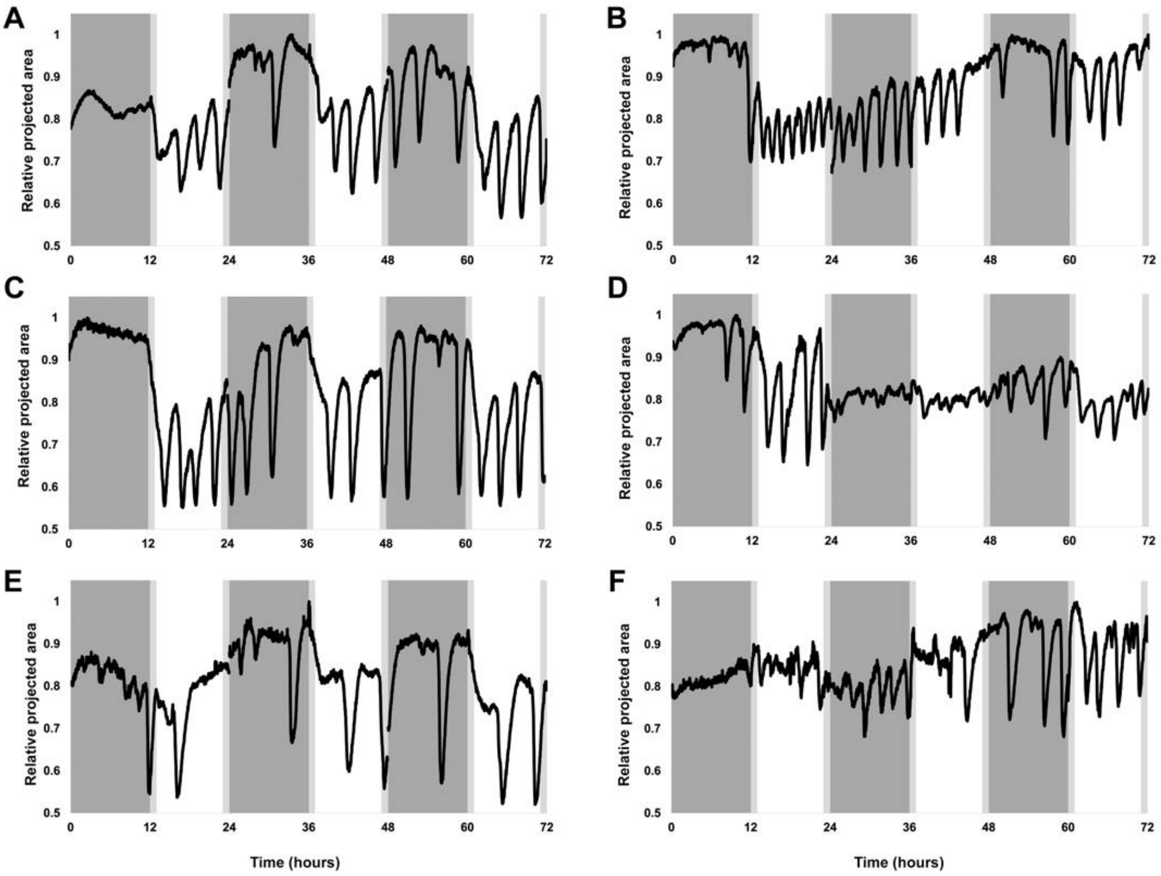


**Fig. S2. Contraction-expansion diagrams of six specimens of *T. wilhelma* (N=6) during the dark/light (DL) experiment.** Contractions are shown as relative changes in projected area. Dark grey indicate periods with total darkness, light grey sunrise/sunset, and white day period.

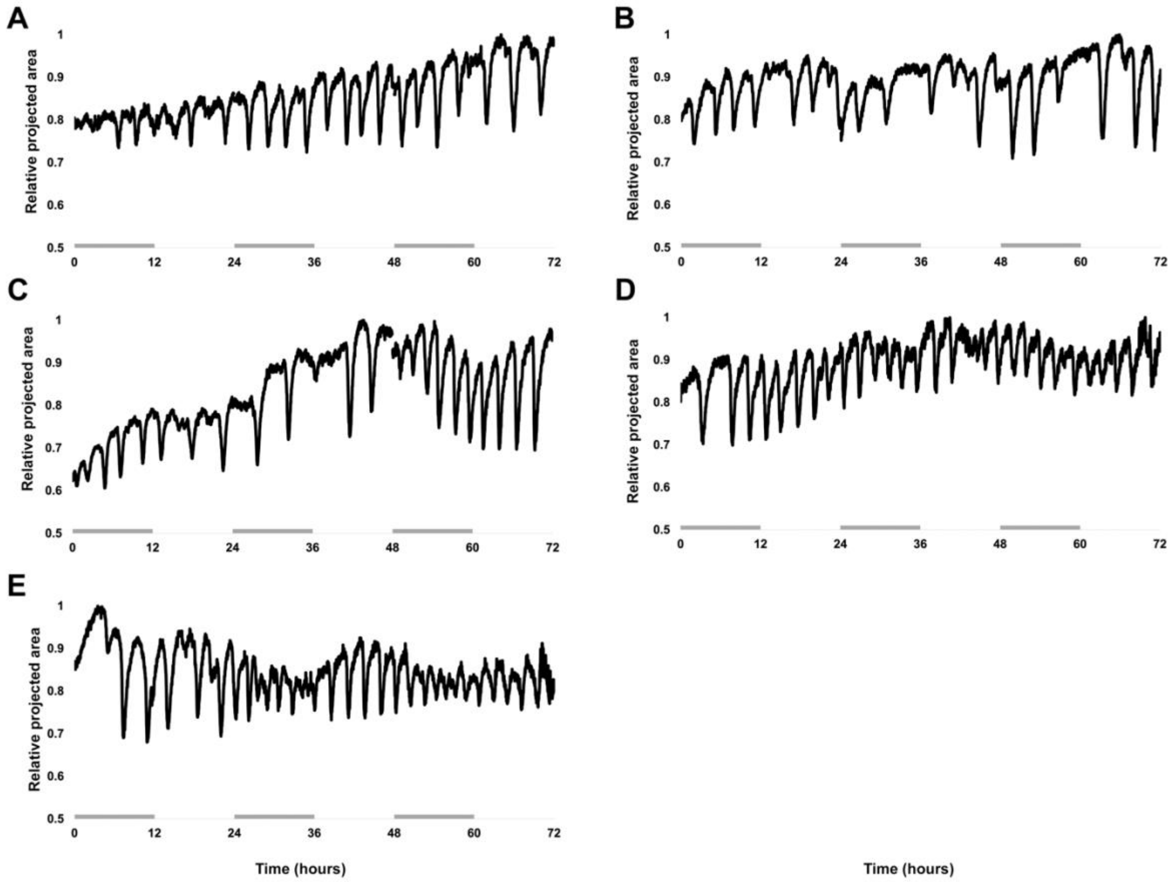


**Fig. S3. Contraction-expansion diagrams of five specimens of *T. wilhelma* (N=5) during the constant dark (DD) experiment.** Contractions are shown as relative changes in projected area. Dark grey bars indicate night periods.

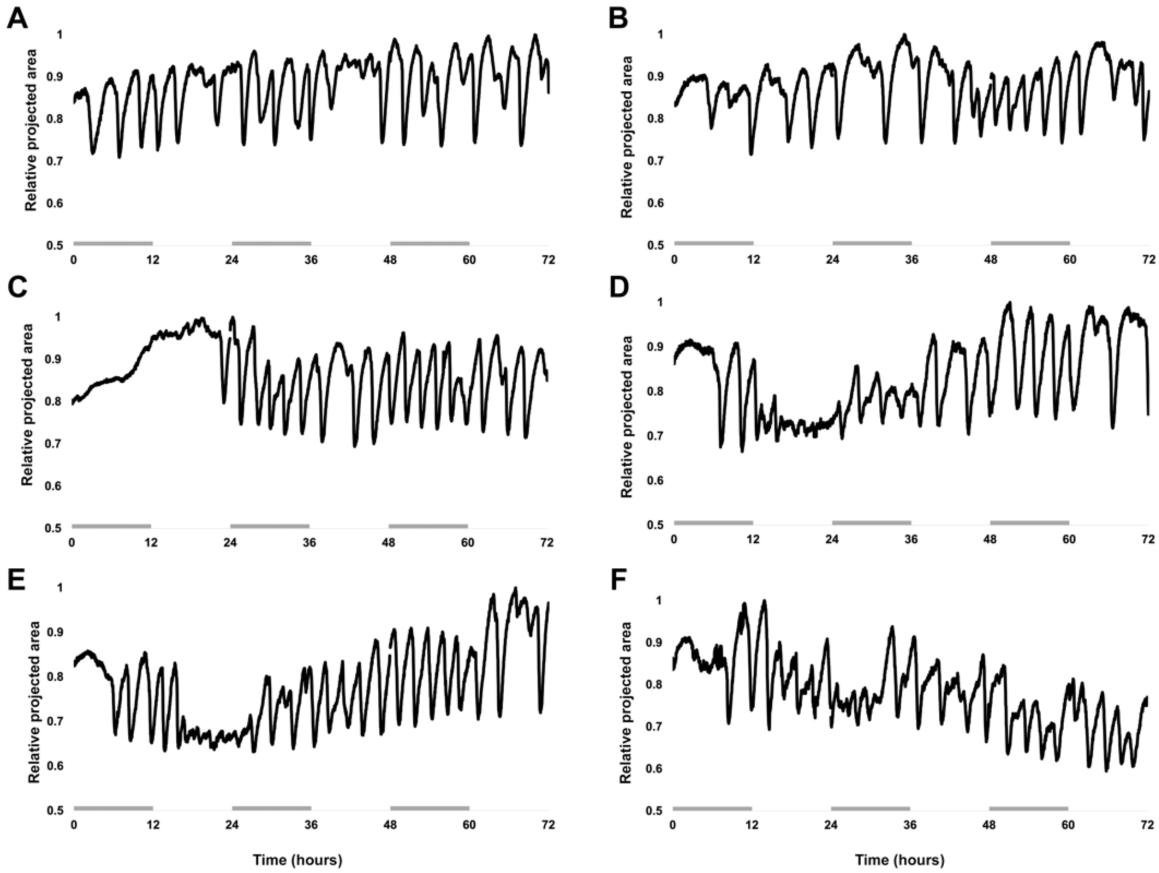


**Fig. S4. Contraction-expansion diagrams of six specimens of *T. wilhelma* (N=6) during the constant light (LL) experiment.** Contractions are shown as relative changes in projected area. Dark grey bars indicate night periods.
